# Cellular responses to proteostasis perturbations reveal non-optimal feedback in chaperone networks

**DOI:** 10.1101/137349

**Authors:** Asmita Ghosh, Abhilash Gangadharan, Sarada Das, Monika Verma, Latika Matai, Debasis Dash, Koyeli Mapa, Kausik Chakraborty

## Abstract

The proteostasis network (PN) comprises a plethora of proteins that are dedicated to aid in protein folding; some with over-lapping functions. Despite this, there are multiple pathophysiological states associated with depletion of chaperones. This is counter-intuitive assuming cells have the ability to re-program transcriptional outputs in accordance with its proteostasic limitations. To this effect, we have used *S. cerevisiae* to understand the route a cell takes as a response when challenged with different proteostasis impairments. Using 14 single deletion strains of genes of Protein Quality Control (PQC) system, we quantify their proteostasis impairment and the transcriptional response. In most cases cellular response was incapable of restoring proteostasis. The response did not activate proteostasis components or pathways that could complement the function of the missing PQC gene. Over-expression of alternate machineries, could restore part of the proteostasis defect in deletion strains. We posit that epistasis guided synthetic biology approaches may be helpful in realizing the true potential of the cellular chaperone machinery.

## Introduction

A nascent polypeptide chain emerges from the ribosome as a linear-chain of amino acids. This polypeptide chain needs to fold into its specific 3-dimensional structure to attain its function. To navigate from an unfolded, linear chain to its specific 3-dimensional structure without misfolding, the cell has a complex network of proteins which assist with either co-translational and or post-translational folding to reach the native structures. Further down the assembly line, other proteins assist in translocation, cellular localization and eventually degradation(1-3). The collective group of chaperones, co-chaperones and degradation proteins that maintain these processes are considered to constitute the Protein Quality Control (PQC) system and are integral nodes in the Proteostasis Network (PN)(4, 5). While proteins are helped by the PQC machinery to attain and maintain their native states, what happens if a PQC component is rendered functionally defective?

Multiple phenotypes associated with pathological disease conditions are known to occur upon chaperone depletions. Mutations in the coding region of a chaperone leading to its loss of function results in a disease phenotype defined as Chaperonopathy(6). Also, in a physiologically more relevant situation, chaperones are depleted in a cell when there is an excessive load of misfolded or aggregated proteins. As an example, Sis1 is known to be sequestered by polyQ aggregates (7) while Aha1 engages extensively with misfolded CFTR Δ508, thus being unavailable to help fold its normal repertoire of substrates(8, 9). We were interested in how the cell responds in a such a scenario of depletion of a particular PQC component. There are three major ways in which the cell may respond to functional depletion of chaperones. First, it can sense the accumulation of misfolded client proteins due to loss of function of the cognate chaperone. In this case the response will be directed by the client proteins which are unable to fold and cell would try to upregulate parallel pathways to take care of the function of these client proteins. Second, proteotoxicity arising due to misfolding or aggregation of substrate proteins can induce transcription of the stress response system to upregulate the entire cohort of chaperone machinery e.g. stress regulated backup chaperones like Ssa3/Ssa4 of Hsp70 and Hsp82 of Hsp90 family respectively in yeast. Third, the cell can sense the loss of specific PQC node and upregulate the existing paralogous machinery of the PQC(10, 11). We still do not understand how the cells respond to depletion of specific nodes of the PQC.

Given the redundancies present in the network of PQC genes and the large number of interconnected components(12), it is expected that the PN can easily adapt to altered demands. This is counter-intuitive when one considers the preponderance of protein-aggregation associated phenotypes and diseases (13). To annotate any specific responses that the cell may launch to tackle loss of a PQC node, we needed to functionally deplete one hub at a time. We chose deletion strains as there isn’t an endogenous way to perturb only one node without affecting other hubs; primarily due to a lack of well-defined substrate exclusivity.

Using a repertoire of 14 PQC gene deletions in *S. cerevisiae* we obtained the transcriptomic adaptation of the cells to these deletions, when different modules of the PN are perturbed. To decipher if re-routing of chaperones (to divide the substrate load) were a limitation during altered proteostasis conditions, we analysed the transcriptomic signature in the PQC deletions. We expected the genes of existing parallel, redundant pathways to be up-regulated as an efficient response to the PQC deletions(14). We obtain a new network of the chaperones based on cellular sensory mechanisms with insignificant overlap with the genetic interaction map; based on epistasis.

Molecules to be deleted were chosen to represent the diverse hubs that govern proteostasis in *S. cerevisiae* (Figure 1A, left panel). Ssa1, as the most canonical and constitutively present Hsp70 while Ssa3 being the inducible Hsp70 protein folding chaperone in the cytosol(10). Hsp82 and its co-chaperone Sti1 to represent the cytosolic Hsp90 system. Hsp104 as a disaggregase and San1 as part of protein degradation. The other classes of proteostatic components were primarily involved in ribosome-associated quality control (Rqc2, Rkr1), ribosome-associated protein folding of nascent chains (Ssb1, Egd1)(15), ribosome assembly (Sse1, Ssb1, Jjj1)(16). Deletion of mcx1 and a hypomorphic allele of ssc1 (DAMP allele) were used to obtain information pertaining to the cellular response due to the loss of mitochondrial chaperones. The abundance of each of these proteins varies (Figure 1A, right panel) under basal conditions in a wildtype strain (17); this provides an insight into housekeeping load that is shared by a particular molecule in the cell.

**Figure 1.**
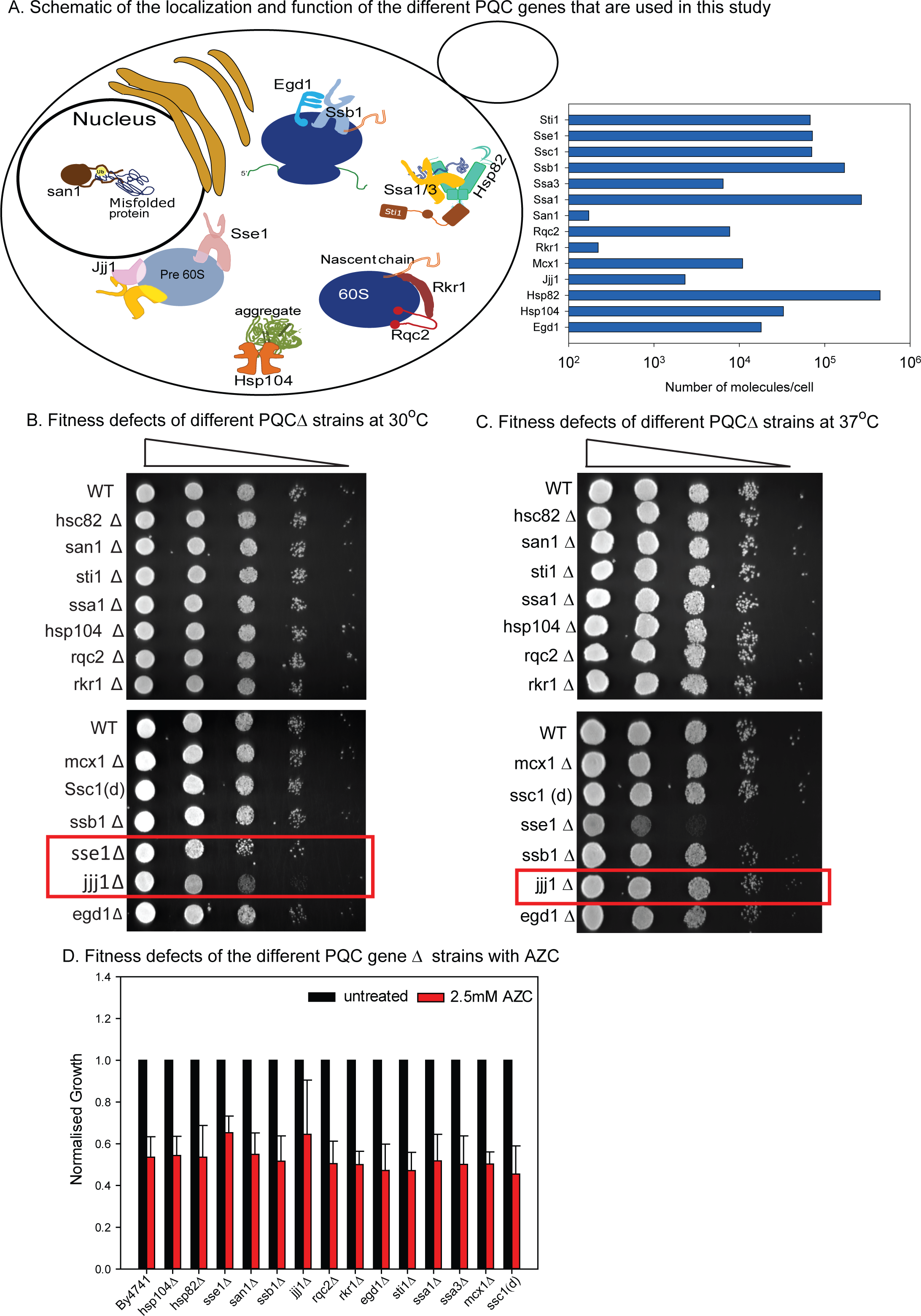
A. Left panel shows localisation and function of each of the PQC genes we picked for the study in the schematic. The right panel shows Number of molecules/ cell for the selected PQC components as reported by Ghammaghemi *et al*.; (17).
B. The deletion (and depletion) strains were spotted on YPD agar plates at 30°C to assay growth fitness of the strains.
C. The deletion (and depletion) strains were spotted on YPD agar plates at 37°C to assay growth defect of the strains upon chronic heat stress.
D. The deletion (and depletion) strains were grown with or without L-AZC at 30°C to assay growth defect of the strains upon chronic misfolding stress.

In this work we show that deletion of protein quality control hubs led to an underlying loss of proteostasis although without any overt fitness defect during normal growth or during proteotoxic stress. Remarkably, it appears that even a phenotypically healthy and exponentially growing cell has limited ability to respond to the specific proteostasis disabilities in the most optimal manner; the redundant machineries were not specifically upregulated as part of the cellular response. We find that activating a parallel pathway, based on epistasis, to over-come the absence of the PQC component benefits the proteostasis capacity of the deletion strain. Thus, it appears that in the chaperone network, cells inherently have a limited scope of response with their transcriptomic responses being decorrelated from their epistasis.

## Results

### PQC deletions have compromised proteostasis

We measured growth of the chosen PQC deletion (and depletion) strains and found only sse1Δ and jjj1Δ to have a defect at 30°C (Figure 1B). Most of the deletion strains did not show a growth defect at 37°C a mild heat shock condition (Figure 1C), or in AZC (Figure 1D) that induces misfolding stress. Surprisingly, that they did not have a drastic impact in cellular fitness even during misfolding stress indicated that either the deleted chaperones played only a minor role, or cellular readjustment nearly perfectly solved the problem arising from the deletion.

To check if cellular readjustment restored proteostasis in the deletion strains, we required a sensor of cellular proteostasis. We designed this using the Nourseothricin Acetyl Transferase gene (NAT-R). We made several temperature sensitive (TS) mutants of this protein in *S. cerevisiae* by screening for resistance to CloNAT (an antifungal/antibiotic) at 37°C and 30°C (Figure 2A). TS-22 was a strong temperature sensitive mutant of NAT-R (Figure 2B and 2C) and degraded faster than WT protein even at 30°C (Figure 2D, S1A). Although mere presence of this mutant did not confer any growth defect to the cells (Figure S1B). This mutant was indeed sensitive to proteostasis as the steady state protein level decreased in the presence of heat shock inducing compound, celastrol, that changes the chaperone pool and hence proteostasis; WT protein was not similarly affected (Figure 2E).

**Figure 2.**
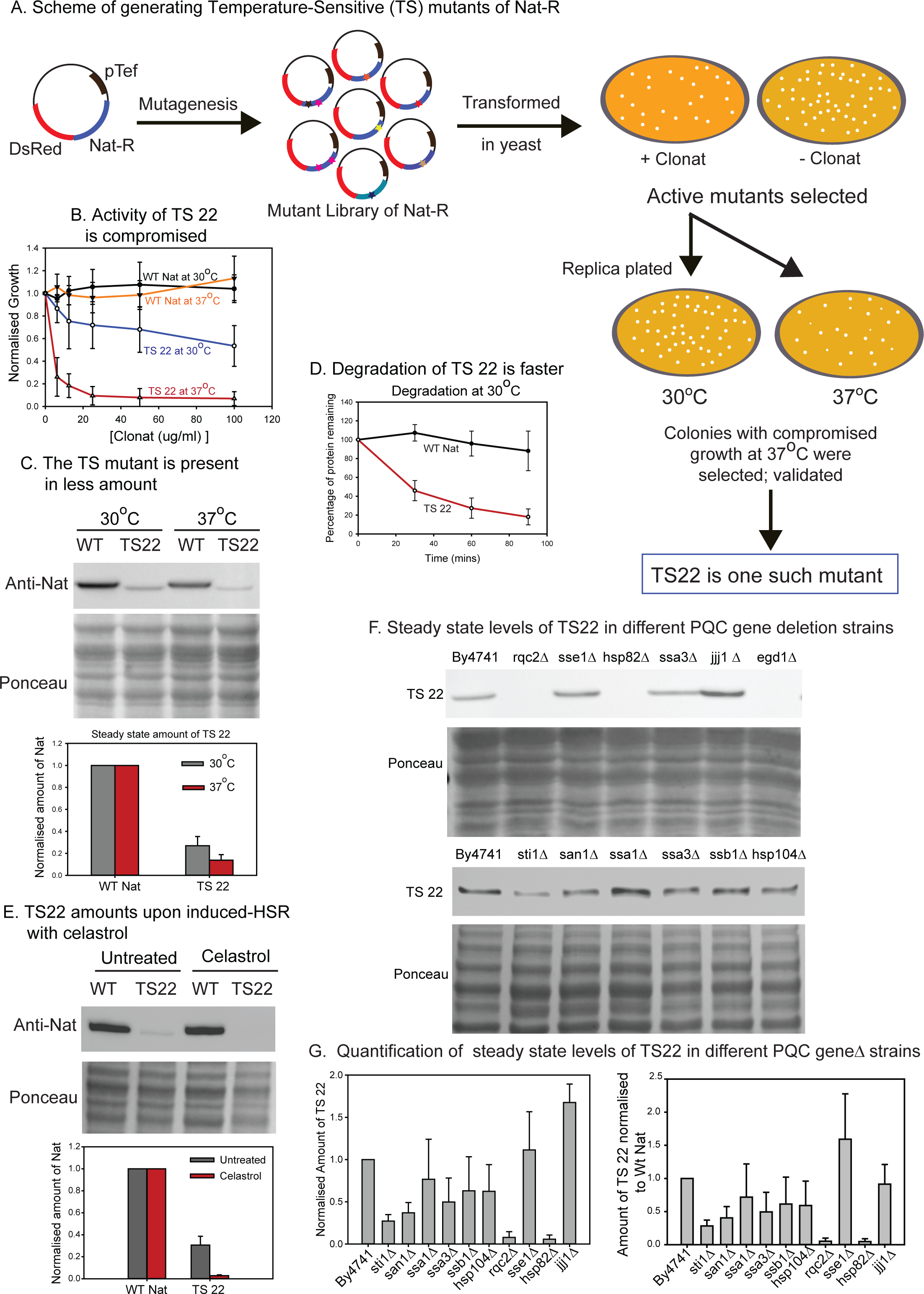
A. Schematic showing the generation of TS mutants of Nat. Multiple mutants were isolated but, only TS22 was used for this study as a candidate metastable protein.
B. Growth assay of Wt Nat and TS22 with varying concentrations of ClonNat at 30°C and 37°C to assay for the functionality of the proteins *in vivo.*
C. Steady state amounts of Wt Nat and TS22 at 30°C and 37°C were determined by western blot analysis.
D. The stability of Wt Nat and TS22 were determined following a cycloheximide chase at 30°C. Ponceau was used for normalisation of protein per lane (loading control).
E. Steady state amounts of Wt Nat and TS22 upon 20μM celastrol treatment at 30°C were determined by western blot analysis.
F. Representative blots for steady state amounts of TS22 at 30°C in different PQC gene deletion strains were determined by western blot analysis. Ponceau was used for loading normalisation.
G. Quantification from 3 biological replicates are plotted for TS22 (left panel). Amounts of TS22 normalised to the amount of WT Nat in the respective chaperone deletion strains are quantitated (right panel).

We used this sensor to monitor the state of proteostasis in the deletion strains (Figure 2F, 2G and Figure S1C, S1D) by measuring the steady-state amount of TS22 with respect to WT protein expression in the same strains. jjj1Δ was special as it exhibited increase in both WT and TS22 levels, indicating an increase in transcription or translation of the gene product. Many of the other strains showed specific difference in the mutant protein indicating differences in proteostasis; these strains mostly did not show any other phenotype either at 30°C or at 37°C or in AZC. This suggested that although the deletions tolerate misfolding stress as well as the WT strain, deleting these genes compromised proteostasis. These results led us to ask how cells respond to these deletions, and why is this not sufficient to restore proteostasis.

### Response to PN pertubations is independent of Heat Shock Response

Towards this we asked if the cells sensed the PQC deletions and mounted any response. Since most of the chosen PQCs were cytosolic, we checked for the reported effect of these deletions on the heat shock response-the canonical cytosolic response(18). Analysis of the published result show that some of the strains like sse1Δ, hsp104Δ, sti1Δ and san1Δ show significant upregulation of HSR reporter (Figure S2A). This was unable to explain why sse1Δ - that induce HSR and hsp82Δ - that do not induce HSR, both show compromised proteostasis. To obtain a global view of cellular response we carried out RNA-seq based transcriptomics for the 14 PQC deletion strains in duplicates (Table S1 and S2). Unlike the synthetic HSR reporter, the expression levels of the canonical Hsf1-dependent HSR-gene ssa4 was high in only sse1Δ. But the expression level of hsp12, a Msn2/4 reporter, was high in most of the PQC deletions (Figure S2B). This categorically negates the possibility that the deletions did not alter cellular homeostasis. There is indication of stress upon depletion of each of the PQCs, the cells sense this and upregulate Msn2/4 pathway and in the case of sse1Δ, the Hsf1 pathway.

Although there is indication of altered proteostasis and cellular response to stress, it was surprising that deletion of abundant chaperones like Ssa1 or Hsp82 did not induce the expression of ssa4. Hsf1 regulates ssa4 expression, and Hsf1 is thought to be the canonical sensor for altered proteostasis in eukaryotes. Importantly, the canonical HSR genes are not enriched in the set of upregulated genes in the PQC deletions (Figure S2C). While the HSR genes were not downregulated in these PQC perturbation strains, the lack of a strong HSR indicates (Table S3) that the general response to PQC deletions do not induce a canonical HSR in *S. cerevisiae* as might be expected. Surprisingly, none of the strains show a significant increase in overall chaperone levels (Table S2).

We asked if the magnitude of response to deletion of a PQC gene is correlated to the expected load on that PQC nodes. The load can be approximated by the physiological protein concentration of each of the PQCs. To obtain the total magnitude of cellular perturbation upon deletion of a PQC node, we took two approaches. First, we measured the total number of differentially regulated genes when a PQC is deleted. Second, we measured the divergence of the transcriptome of a PQC deletion strain with respect to WT cells (Table S4). Both these measures of magnitude of response did not correlate with the copy number of the deleted PQC protein (Figure 3A and 3B). For example, protein copy of Ssa1 is higher than Sse1, yet deletion of sse1 causes a larger transcriptome reorganization than ssa1 deletion.

**Figure 3.**
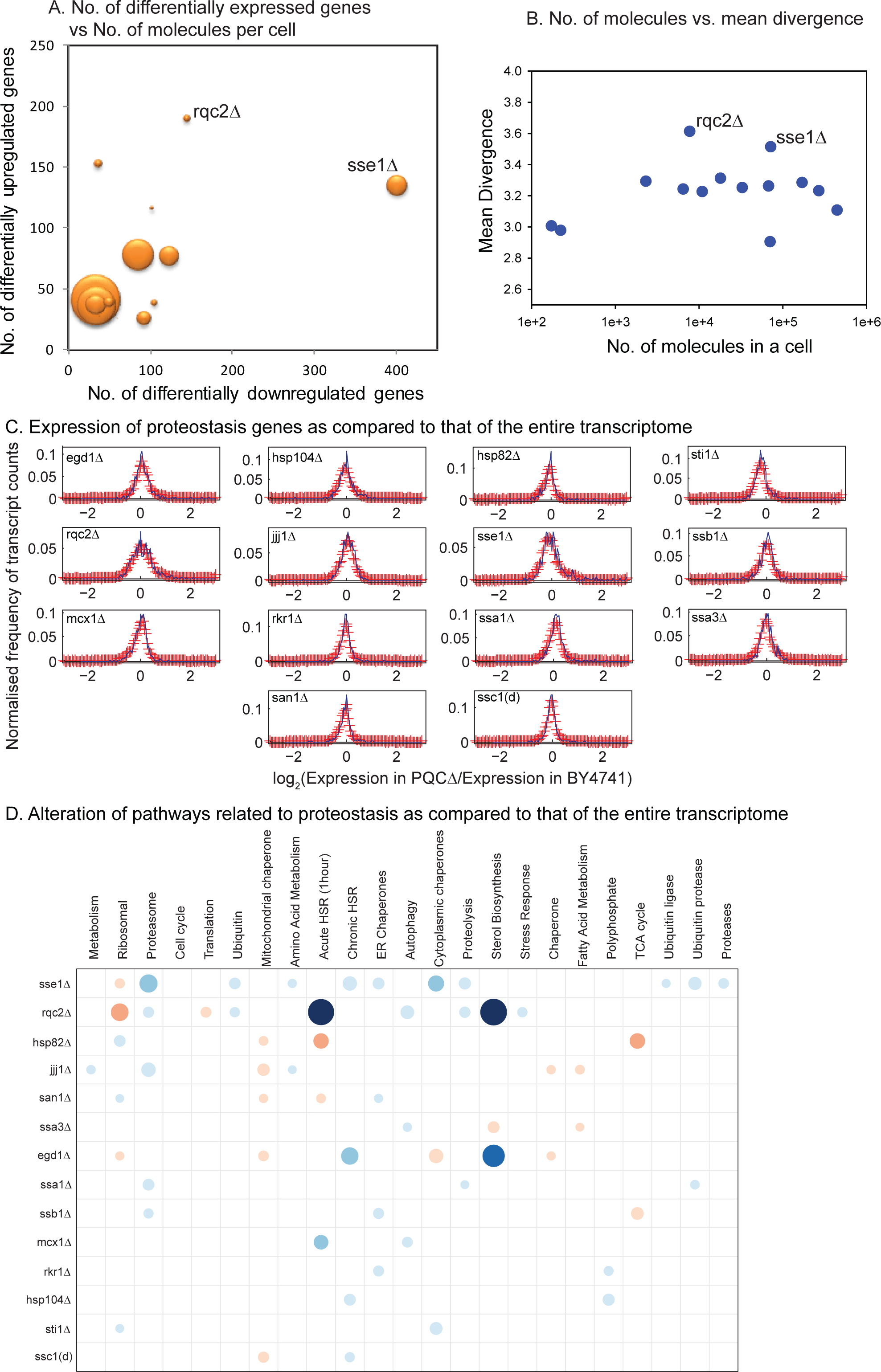
A. Number of differentially upregulated and downregulated genes in the PQC gene deletions were plotted against the number of protein molecules of each PQC (as obtained from (17)).
B. Mean divergence is plotted against the number of protein molecules of each PQC component.
C. We calculated fold change of all the transcripts with respect to BY4741 for each of the deletion strains. The proteostasis component genes were defined as in Table S4. In all the deletion strains, proteostasis network set have lower divergence than a randomly chosen subset of the same number of genes. For each case, to calculate Shannon Entropy we obtained a normalized histogram with 121 bins ranging from −3 to 3 of log_2_ of fold change values. The area of the histogram was normalized to 1, in case the distribution is more spread out, the divergence of the distribution is high. (Also see Table S4, S5).

The cellular response to PQC gene depletion could be initiated by loss-of-function of clients, or by proteotoxicity arising from misfolded proteins. Aiming to segregate the effects, we characterized the responses in terms of proteostasis components that are differentially expressed in these PQC deletions. The divergence of proteostasis components from the WT cells were generally low compared to the rest of the genes (Figure 3C and Table S4, S5) indicating that proteostasis components were on an average less perturbed than other cellular components. Proteasome/protein clearance pathways are the only ones related to proteostasis that are consistently, but mildly, upregulated in many strains (Figure 3D, Table S6). This indicated that the response was most probably dominated by the effect of non-functional client proteins and not due to protein aggregation or misfolding. This corroborates nicely with the absence of Hsf1 dependent signal. This also explains the loss of proteostasis as measured by the sensor; cellular response was directed towards taking care of pathways lost due to client-misfolding and not proteostasis defects. Interestingly, jjj1Δ and sse1Δ show the highest perturbation in proteostasis genes according to this measure suggesting that these strains have the largest proteotoxicity dependent response; these strains are also the only ones that show a measurable growth defect.

Taken together, each of the PQC deletions reorganize their transcriptome sensing some stress; the reorganization is however not dependent upon the canonical heat shock pathway. This and other evidences indicate that the primary stress in these deletion strains arise due to loss of function of clients and not due to misfolding. However, there is a mild misfolding stress that in many cases is dependent on the Msn2/4 axis, but overall, the response is not correlated to the expected load on the deleted node of PQC.

### Basis of cellular response to chaperone deletion

If the response primarily is the result of non-folding of clients, deletion of chaperones that have overlapping substrates should up- or down-regulate similar genes. In order to check similarities between the responses of two different deletion strains, we obtained the pair-wise correlation coefficients between the transcriptome profiles of different chaperone deletion strains (Figure S3A). This was done by plotting the correlation between the transcriptome profiles (fold-change values of a chaperone deletion with respect to WT). The strength of the correlations indicates similarity between the responses of two chaperone deletions. A hierarchical clustering of the correlation matrix (Figure 4A) also depicted similarities in the transcriptomic response that was apparent within multiple chaperones. Hierarchical clustering identified four main clusters in the plot. First, a cluster of two cytosolic (Jjj1, Ssa1), and two mitochondrial (Mcx1, Ssc1) PQC nodes were formed (cluster 1). Second, the more heterogeneous group consisting of chaperones (Sti1, Hsp82, Ssb1) along with two E3 ubiquitin ligases (Rkr1, and San1) (Cluster 2). Co-clustering of San1 and Rkr1 along with other chaperones, highlight the importance of these two proteins in proteostasis maintenance in general. Third cluster was found to contain Egd1 and Rqc2 (Cluster 3), both of which are associated with ribosomes and are involved in quality control of nascent polypeptides. Interestingly, Rkr1 (Ltn1), known to be associated with Rqc2 (Tae2) was not part of this cluster. Fourth, cytosolic chaperones that aid folding and refolding of aggregates (Hsp104, Ssa3 and Sse1) clustered together (Cluster 4).

**Figure 4.**
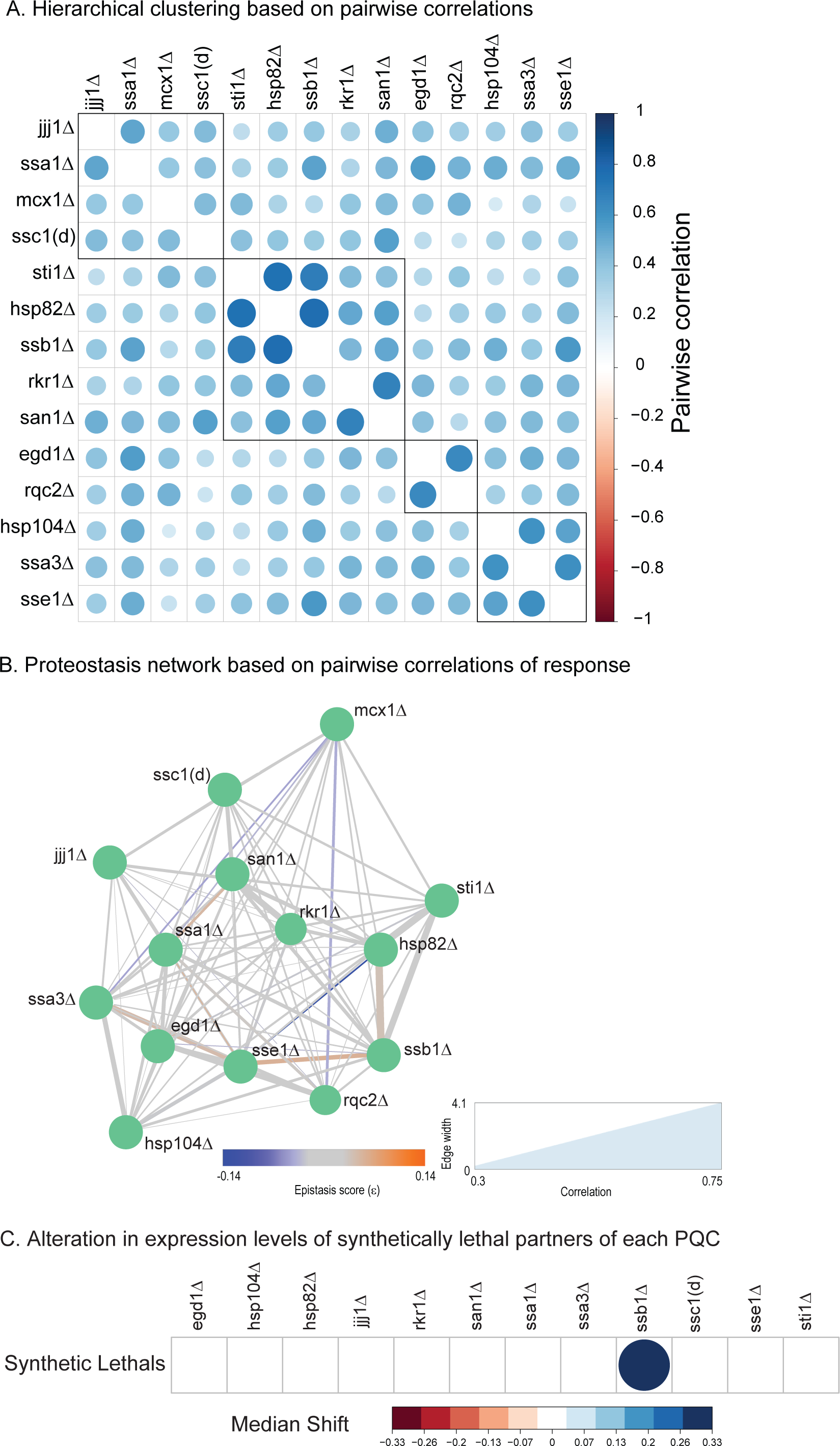
A. Hierarchical clustering for PQC genes using the correlations obtained in (S3A). Four clusters are identified and are indicated by black boxes. Bigger circle indicates a stronger correlation. The colours and size of the circle indicate the extent of correlation between the transcriptome profile of two gene deletions strains. The diagonal elements have a correlation of 1 and are omitted for clarity.
B. The network view of the transcriptomic correlations between PQC deletions. A thicker line indicates stronger correlation than a thinner line. The colour of the lines are dictated by the Boone epistasis scores (ε) (20) between the two genes. The 3- colour key for epistasis showing blue for negative epistasis (exacerbating interactions), grey indicates zero epistasis (no interaction) and red shows positive epistasis (alleviating interactions).
C. Expression changes of synthetic lethals in each of these PQC gene deletions were compared to that of the entire transcriptome. The fold change in their expression values in the PQC deletion strains w.r.t By4741 (parental strain) was compared to the global alteration in transcripts using Mann-Whitney test. The p-values obtained were corrected for multiple comparisons using FDR and the cells that did not cross the cut-off significance for p-value 0.05 are without any colour. The colour scheme is graded from red (down-regulated) to blue (upregulated) showing median fold change values. The size of the circle represents the median fold change shift in either direction.

These correlations were used to obtain a network between the PQCs based on the similarity of transcriptomic response. We overlayed the transcriptomic response network with the epistatic scores between the chaperones and obtained an unexpected dissimilarity between the epistasis scores and response correlations (Figure 4B). With respect to epistasis, genes in the same pathways are expected to have positive epistatic scores. These pairs, since they are on the same pathways, should also show similar cellular response upon their deletion. However, the positively epistatic pairs do not show a stronger correlation than the negatively epistatic pairs. Strikingly, although Sse1 and Sti1 are strongly exacerbating pairs in terms of epistasis (negative epistatic interaction), they have one of the weakest links in the map. Additionally, many of the non-epistatic pairs (for example San1 and Rkr1) show the strongest connections in the response map.

Overall there seems to be a weak similarity between the PQC nodes based on their localization and function. This corroborates well with client-dependent response rather than misfolding/aggregation driven one. Despite this, similarities highlight interesting overlaps that are yet to be discovered: connection between Ssb1 and Hsp82, or Rqc2 and Egd1 will be interesting to delineate.

Yeast PQCs are heavily redundant due to genome duplication and are known to have functional redundancies(10, 19). We asked if the response to PQC deletion led to upregulation of the parallel pathways(14). To answer this, we investigated the correlation between reported epistasis values (ε) and gene expression(20). Genetic interactions as obtained through fitness epistasis scores (ε) define functional relationships between pathways; their dependency, and redundancy. Genes with strong negative interactions are usually the ones that operate in parallel although redundant pathway; we expect the pathways parallel to a specific PQC-containing pathway to be upregulated when that PQC is deleted. Given this, we wondered if the cellular response is guided by the genetic interaction network of the PQC genes. We addressed the issue by checking for the correlation between epistasis scores (ε) and the transcriptional response to a PQC deletion and see if the response is guided by the epistatic interactions defined by the chaperone (Figure S3B). However, we observed no apparent correlation between epistasis scores (ε) and the transcriptome regulation. Thus, the experimentally observed redundant pathways are not upregulated upon deletion of a PQC. The strongest negative genetic interactors would be synthetically lethal; even these genes were not upregulated (Figure 4C). Summarily, the backup pathways, or the parallel pathways are not upregulated when a particular PQC is deleted.

Taken together all of the above results support that the response to PQC gene deletions are guided by missing function of the chaperone-clients in the cell and not through sensors of proteostasis. This response possibly compensates for the loss of activity of the clients but not for the loss of proteostasis. This seems to be a fundamentally different from the heat shock response.

To verify if this limitation was specific to PQC gene deletions or true for any gene deletion in *S. cerevisiae,* we took advantage of a recently reported large scale transcriptional study of many deletion strains(21). This allowed us to investigate if transcriptional response to specific deletions is generally uncoupled to epistatic interaction scores or if this feature was exclusive of the PQC genes. None of the gene deletions, except three (ybl039c, yor209c, ygl252c), show any significant correlation between the transcriptional alterations of the expressed genes and the epistasis of these with the gene that is deleted (Table S7). All the three that show significant correlation exhibit negative correlation between epistasis values and expression change; negatively epistatic genes are upregulated while the positive ones are downregulated. This is exactly as we expected in case optimality is built into the response network. Thus, barring these three exceptions, all the other deletions behave like the PQC deletions; genes that are negatively epistatic to the deleted genes, do not show an upregulation. This underlines that response to any perturbation, not limited to the PN, is not guided by the optimality as predicted by the genetic interaction network

### Cryptic optimal pathways can be activated to alleviate PQC deletion associated fitness defects

We wondered would the cellular growth defect due to deletion of a PQC may be relieved in case there was an optimal response. Two PQC gene deletions (jjj1Δ and sse1Δ) showed significant growth defect at physiological temperature of 30°C (Figure 1B) and also the strongest perturbation of proteostasis pathways (Figure 3, Table S4). As a test to check if forced upregulation of a parallel pathway can alleviate a PQC deletion we overexpressed Sse2, a negative genetic interactor of Sse1, in the absence and presence of Sse1. Sse2 overexpression indeed alleviated the growth defect of sse1Δ deletion strain (Figure 5A and S4A). This demonstrates that cryptic pathways exist, but these are not upregulated optimally to completely take care of the PQC deletion.

**Figure 5.**
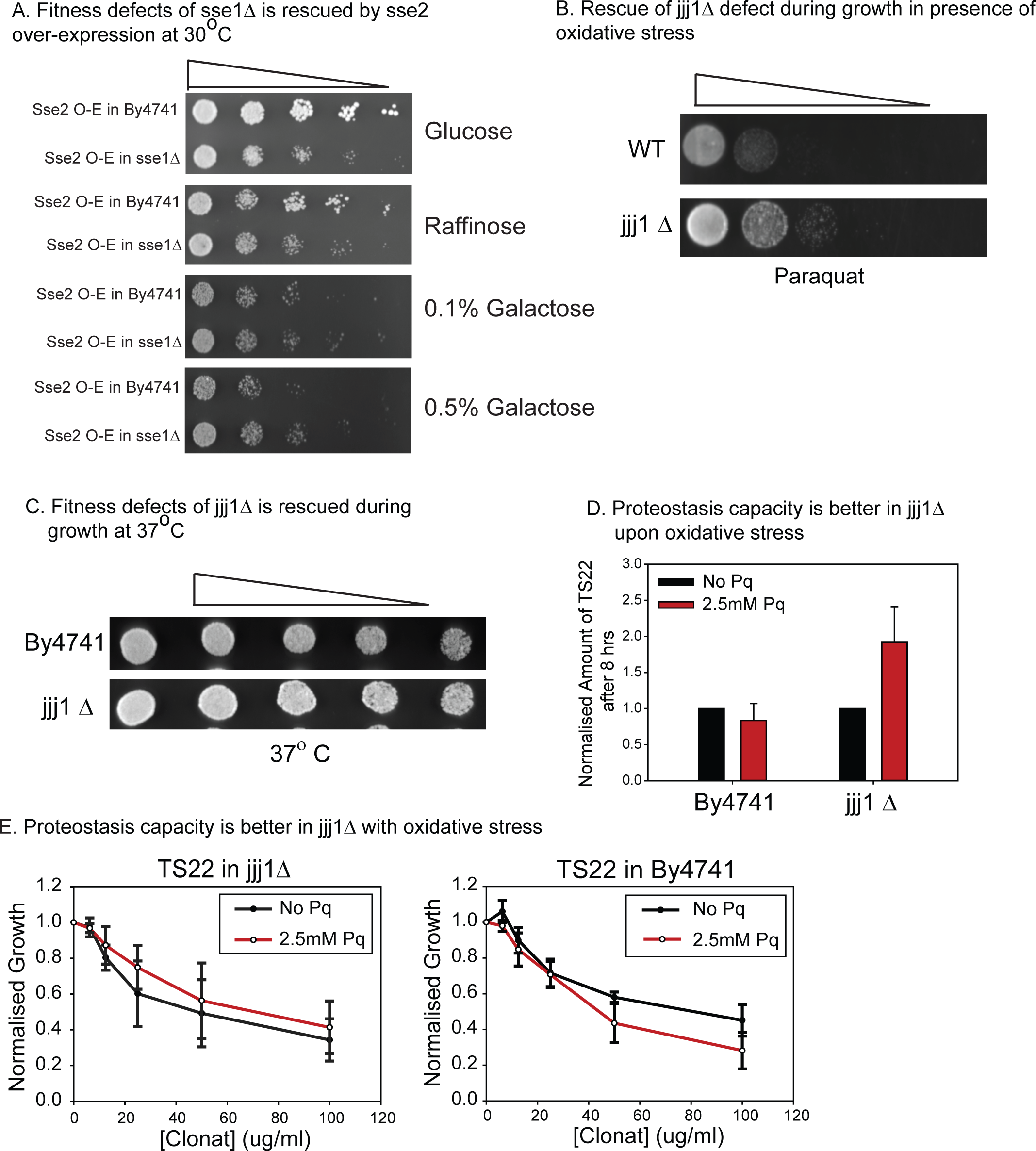
A. By4741 and sse1Δ bearing galactose-inducible over expression of Sse2 were spotted on YPD, YP + Raffinose or YP+ Galactose and grown at 30°C.
B. By4741 and jjj1Δ were spotted on YPD agar plates containing 5mM Paraquat and grown at 30°C to assay growth fitness of the strains.
C. By4741 and jjj1Δ were spotted on YPD agar plates and grown at 37°C.
D. Quantification of western Blotting of TS22 in By4741 and jjj1Δ with or without 2.5mM Paraquat (refer S4C).
E. Activity of TS22 measured in terms of growth in varying concentrations of the antibiotic CloNat, in jjj1Δ and By4741 with or without 2.5mM Paraquat.

However, we did not get fitness alleviation in jjj1Δ when we expressed 3 of the randomly chosen top negative genetic interactors (asc1, vid22, and vam7) (Figure S4B). Since all the four proteins work as part of some complex in a pathway, it is apparent that upregulation of only a single gene is not enough to upregulate the cryptic pathway. One of the top negative interactors of jjj1Δ is sbt5 (22) and this encodes a transcription factor that is activated during oxidative stress. Remarkably, activation of this branch by treating the cells with a mild concentration of paraquat (Pq), could alleviate the growth phenotype of jjj1Δ (Figure 4B). Similarly, a mild but chronic heat shock treatment (at 37°C) that is also known to upregulate oxidative stress response pathways, also alleviated the growth defect of jjj1Δ (Figure 4C). Thus, cryptic pathways could be activated to rescue defects due to the tested PQC deletions. This also demonstrates that fitness compensation is dependent upon activation of redundant pathways rather than a single gene upregulation.

To further check if proteostasis defect of jjj1Δ is also alleviated by upregulating the oxidative stress response pathway, we monitored the stability of the metastable TS22 protein upon paraquat treatment. Assuringly, TS22 was more stable in jjj1Δ in the presence of oxidative stress than in the absence of the stress (Figure 4D). This was true only for jjj1Δ and not WT-strain where the protein was destabilized in the presence of oxidative stress. Commensurately, the activity of TS22, but not WT-NAT, was marginally higher in jjj1Δ in the presence of Pq than in its absence (Figure 4E). Oxidative stress response pathway, that shows interaction at the genetic level, could be activated to compensate for the absence of jjj1. However, a naturally evolved cell has not linked these two pathways to compensate for the absence. Taken together, the above examples demonstrate that cellular systems do have the capability to take care of the short-comings, but this capability is not switched on by default upon depletion of the gene.

### Implications

Proteostasis network, like all other cellular networks, has a complex architecture that needs to be understood to tackle problems that are associated with its collapse. Proteostasis collapse is a hallmark of ageing and is also associated with late-onset diseases(23, 24); alleviation of its collapse has the potential to delay progressive ageing and age-associated diseases. Active research in this field focuses on using the cellular response pathways, such as the HSF1-mediated heat shock response pathway, to revive proteostasis (25, 26). The PQC component deletions used in this study were found to have an underlying defect in proteostasis capacity. Despite this, we uncover that genes part of the proteostatic network are only mildly altered, which is far less than the alteration of other cellular pathways upon a PQC component deletion/depletion. This is surprising and our current work shows that deletion/depletion of functionality of certain nodes may not be reversed by harnessing the existing stress response pathways. These canonical stress response pathways are inherently limited in scope as they are not coupled to the functionality of the nodes. This limits the cellular ability to restore proteostasis and may be one of the reasons why cells are unable to handle aging problems. Although the work reported here is done on *S. cerevisiae,* and with single deletions of chaperones, it is likely to hold true for more complex organizational structures in mammals. It is also likely that the response pattern and efficiency is dependent on cell and tissue-type dependent on the background response pathways. Only further work in this direction will be able to substantiate the findings and their generality. Since this work is based on PQC gene deletions and not partial depletion it may not correctly represent scenarios where chaperones are sequestered. However, deletions represent the extreme limit of depletion of a protein, it is therefore unlikely that cellular response dormant even after deletion will be switched on upon partial depletion of the protein. Summarily, this study unravels a cryptic limitation in the yeast chaperone network, a proposal that needs further investigation in higher organisms.

## Materials and Methods

### Media and growth conditions

Rich media for culturing yeast (YPD medium) containing 1% (w/v) yeast extract, 2% (w/v) peptone and 2% (w/v) dextrose. Yeast strains BY4741(leu2Δ0 ura3Δ0 met15Δ0 his3Δ1) and deletion strains in the background of BY4741 *(Saccharomyces* Genome Deletion Project, http://www-sequence.stanford.edu/group/yeastdeletionproject/deletions3.html) (Open biosystems, GE Healthcare) were grown at 30°C, 200 rpm.

Cells were innoculated in YPD at 0.2 O.D. 600 from an overnight primary culture and grown to mid-log phase before harvesting for RNA isolation.

RNA sequencing for each strain was performed in two biological replicates.

### Growth phenotype by spotting

From overnight primary culture in YPD, cells were innoculated in YPD at 0.1 O.D. and allowed to grow at 30°C, 200 rpm they reached an O.D. of 0.7-0.8 thereafter, they were serially diluted (Starting with an O.D. of 0.5) by a factor of 10 and 5 μl of each dilution was spotted on YPD plates that either contained a stressor of did not. These plates were then kept at 30°C for 48 hours; except in the case of heat shock, they were incubated at 37°C. Over-Expression Plasmids for Sse1, Sse2, Vid22, Vam7 (27) were obtained from Dharmacon.

### Western Blotting

Secondary culture was innoculated at ~ 0.1 O.D. and grown at 30°C; 200rpm. After 6 hours of growth, the cultures were pelleted. Protein was isolated by alkaline lysis method, concentration was estimated by BCA (Thermo Scientific, USA) and 30μg total protein was loaded per lane in a 10% gel. The nitrocellulose membrane was probed with custom anti-Nat-r primary antibody (1: 5000 dilution). A horseradish peroxidase-conjugated goat anti-rabbit antibody (sc-2030, Santa Cruz Biotechnology, Inc. USA) was used as secondary antibody and developed through chemiluminescence (Immobilon Western Chemiluminescent HRP Substrate, Merck Millipore). Blots were quantitated using ImageJ and were normalized to ponceau with its respective blot.

### Growth Assay

From an overnight primary culture, cells were grown in 96-well, deep-well plates for the assay with varying concentrations of the antibiotic CloNat (nourseothricin; AB102XL from

Lexy NTC). 400ul of culture (initial O.D. being 0.05) was kept per well and grown at 30°C; 200rpm for 12 hours. Growth readings were taken on a multi-plate reader (TECAN).

### RNA preparation

Cells were lysed using acid-washed glass bead and total RNA was extracted by TRIzol (Invitrogen) method. It was further purified using Qiagen columns (RNeasy mini kit). RNA concentration and integrity were determined using nanodrop and agarose gel electrophoresis.

### Library preparation

Using Truseq RNA sample prep kit v2 (stranded mRNA LT kit), the library was prepared according to the manufacturer’s instructions (http://www.illumina.com). 700ng of RNA from each sample was used to generate the library. Adaptor-ligated fragments were purified using AMPure XP beads (Agencourt). Adaptor-ligated RNAseq libraries were amplified (12-14 cycles), purified and measured with a Qubit instrument (Invitrogen). The average fragment size of the libraries was determined with a BioAnalyzer DNA1000 LabChip (Agilent Technologies). Diluted libraries were multiplex-sequenced and run on a HiSeq 2000 Illumina platform in HiSeq Flow Cell v3 (Illumina Inc., USA) using TruSeq SBS Kit v3 (Illumina Inc., USA) for cluster generation as per manufacturer’s protocol.

### Mapping sequence reads

Only reads with phred quality score equal or higher than 30 were taken for analysis. Trimmomatic (v0.43)(28) was used to trim read sequences. Reads were then aligned to the transcriptome of *S. cerevisiae* strain S288c as available from ENSEMBL using Kallisto (v0.36) software(29). The reads on an average had over 80% alignment with the reference genome. Gene expression levels were estimated as TPM (Transcripts per million) values. The estimated counts were combined into a matrix and analyzed with EBSeq v1.10.0. Differential expression tests are run using EBTest with 20 iterations of the EM algorithm(30). After this, the list of differentially expressed genes, log_2_fold change of all the genes in the transcriptome and the posterior probabilities of being differentially expressed are obtained using GetDEResults with method “robust”, FDR method “soft” FDR = 0.05, Threshold FC 0.7, Threshold FC Ratio = 0.3.

### Construction of the Co-regulation Network and hierarchically clustered corrgrams

To examine the similarity of the transcriptomic response to chaperone deletions, we computed the pair-wise correlation co-efficient of log_2_ fold changes of all the genes (complete transcriptome) across all the strains. Here, the correlation co-efficient indicates the extent of similarity of the transcriptomic response across mutants to the loss of the chaperone (1 being most similar and 0 being least similar). The correlation matrix was converted into a network with nodes representing the mutants and the edges weighted by the correlation between the mutants. A force directed layout (implemented through Cytoscape(31)) was then applied on the network which clustered strongly connected nodes towards the centre and repelled weakly connected nodes to the periphery of the network. The nodes of the network show the structure of the protein, if available and were taken from STRING database(32). A complementary representation involves hierarchically clustering of a heatmap of the correlation values.

### Protein Complexes and Pathways, expression shift Analysis

To detect shifts in expression for pathways, we employed a GO pathway/functional category based analysis strategy that tests for the overall shift in expression levels for a group of genes that belong to a pathway (obtained from YeastMine) against the rest of the genes in the transcriptome. Refer to Table S4 for details of genes used for each category. For the category 1 hour HSR (Heat Shock Response), we took the first 45 genes upregulated upon 60minute heat shock treatment at 37°C by analysing a previously published microarray data (33). For 6 hours chronic HSR, we took the top 40 genes from our data, upon growing By4741 at 37°C for 6 hours (till mid-log phase). The non-parametric Mann Whitney test is used to avoid issues with over-dispersion in the data and to test the significance of the median shift in expression of genes in a particular pathway in comparison to the whole transcriptome. The Benjamini-Hochberg correction is applied when testing in a combinatorial fashion across pathways to correct for multiple testing.

Besides GO pathways we followed the same strategy to examine the expression status of physically interacting partners, co-chaperone genes, and synthetic lethal partners. The list of physical interacting partners for each protein was obtained from BIOGRID(34). All statistical tests and analysis were performed with R version 3.3.0 (35)scatterplot matrices using the lattice package and heatmaps were made using the corrplot package(36).

### Transcription - Epistasis correlation

We downloaded the transcriptome fold changes as provided in the supplementary files by in a recent report by Kemmeren et al.,(21) and mapped the deletion from the columns in the provided data set to deletions in the epistasis score matrix and extracted their epistasis profiles. 511 mutants out of the 700 mutants mapped to the epistasis score matrix. For each deletion, we computed the correlation co-efficient with significance indicated by P-values between the transcriptome fold-changes of genes to the epistasis score within the epistasis profile.

### Calculating Divergence

We calculated fold change of all the transcripts with respect to BY4741 for each of the deletion strains. The proteostasis component genes were defined as in Table S5. For each deletion strain, from the whole transcript sample we calculated the fold change distribution for 10,000 sets of randomly picked genes (with replacement) to match the number of proteostasis genes (611). As a measure of divergence of the distribution, Shannon Entropy was calculated for each of these 10,000 distributions, the mean of the shannon entropy (A) and the standard deviation (B) were obtained. Shannon entropy was also calculated for the proteostasis components (C). Difference of the Shannon Entropies (D) were obtained as C-A. Negative value indicates lower divergence in proteostasis components. p-values were obtained by using z-test (E). In all the deletion strains proteostasis components have lower divergence than a randomly chosen subset of the same number of genes, and this difference is statistically significant.

Note: For each case, to calculate Shannon Entropy we obtained a normalized histogram with 121 bins ranging from −3 to 3 of log2 of fold change values. The area of the histogram was normalized to 1. The fraction in each histogram was read directly as the prbability of finding genes in a particular (ith) bin (pi). Shannon entropy was obtained by taking a summation of - pi*log(pi) for all the bins. 0 would indicate lowest entropy state, in case all genes were in the same bin. The value will increase in case the distribution is spread out, in other words if the divergence of the distribution is high.

## Acknowledgement

We are grateful to Dr. Mohammed Faruq for aiding us with the Illumina sequencing platform. We thank Dr. Deepak Sharma (affiliated to CSIR-IMTECH) for assistance with reagents. This work was primarily funded by OLP1104 grant by CSIR to KC and partially by SNU core-funding to KM. We thank the HPC facility of CSIR-IGIB, for aiding us with computing resources. AG1, AG2 and LM thank UGC for their fellowship. MV is grateful to CSIR, and SD to SNU-core funding for their fellowship.

## Author Contributions

AG1, KC, KM designed the work. KC, KM and DD supervised the work and analysis. Sequencing was done by AG1. LM made the TS mutants. AG1, SD, MV did the yeast experiments. Analysis was primarily done by AG2 along with AG1 and KC. AG1 and KC wrote the manuscript with input from all authors. All authors read and approved the final version of the manuscript. Authors declare no competing interests.

---

**Figure S1.**
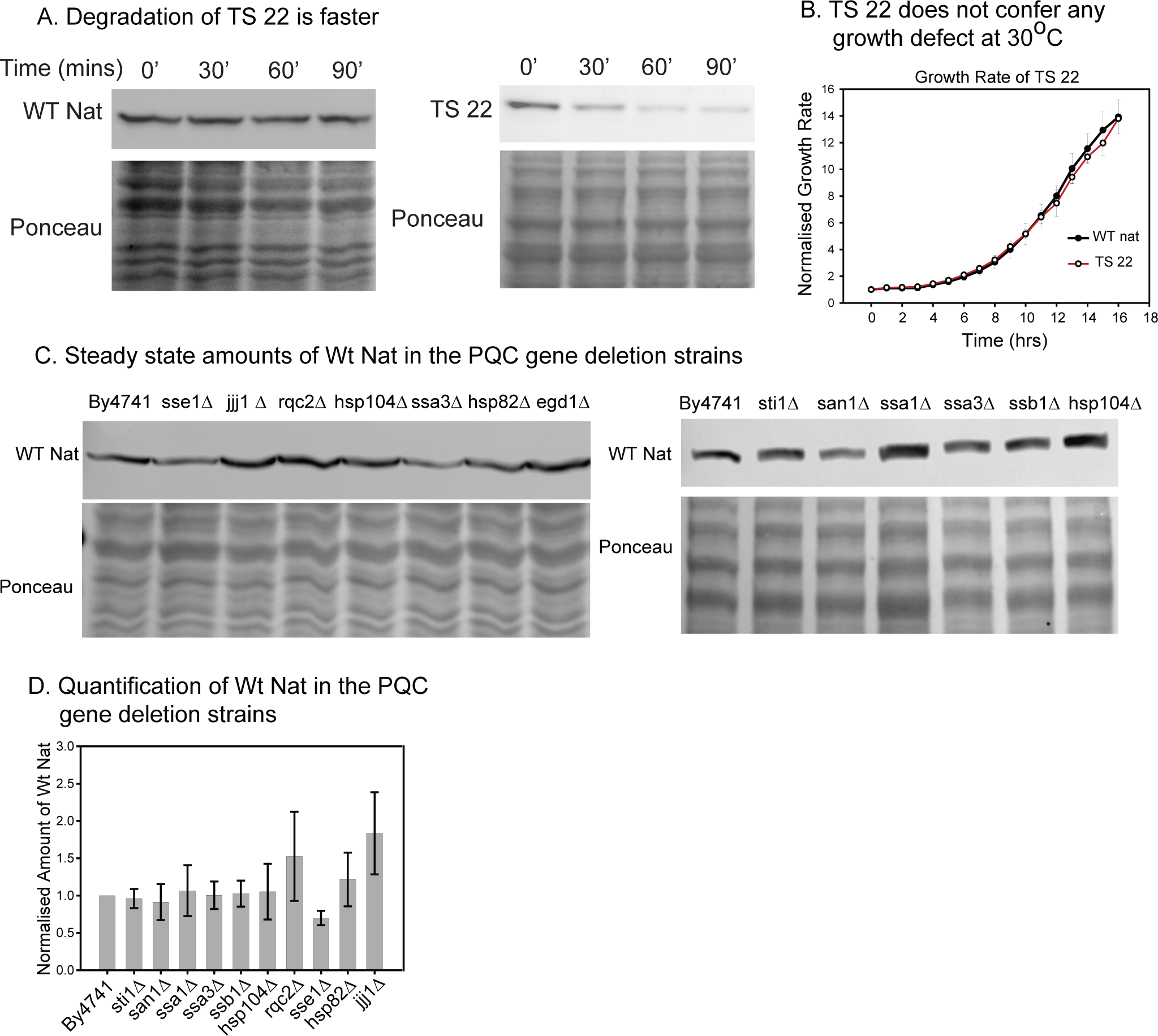
A. Representative blots for cycloheximide chase of TS 22 and WT Nat are shown. Ponceau was used for loading normalization.
B. Growth curve of Wt Nat and TS22 at 30°C done on Bioscreen. Average of 3 biological replicates is plotted.
C. Representative blots for steady state amounts of Wt Nat at 30°C in different chaperone deletion strains were determined by western blot analysis. Ponceau was used for loading normalisation.
D. Quantification from 3 biological replicates are plotted for WT Nat in different PQC gene deletion strains.

**Figure S2.**
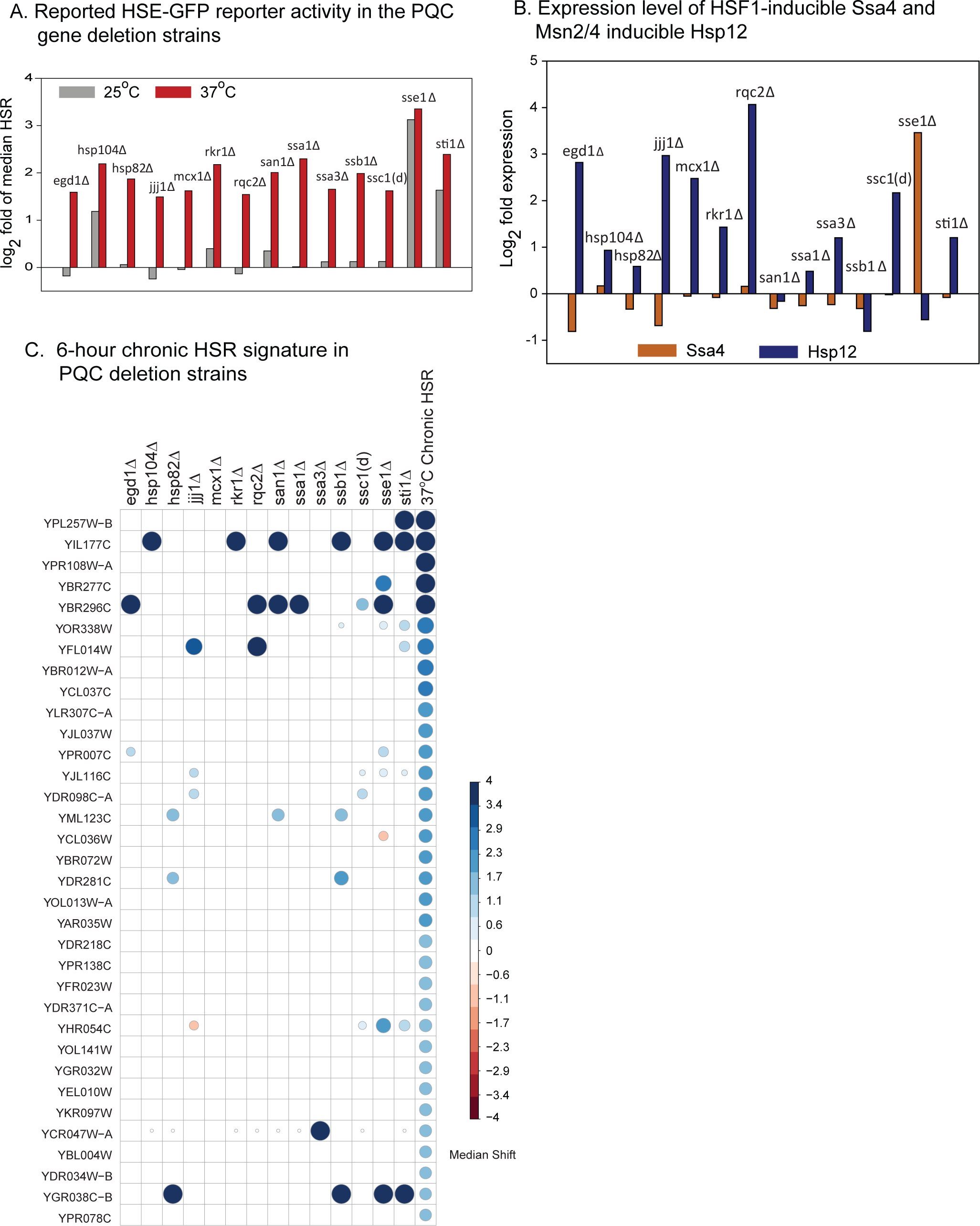
A. HSE-GFP reporter levels in these PQC deletions at 25°C and 37°C as obtained om Brandman *et al*;,(18)
B. Expression levels of HSF1-inducible ssa4 and PKA-inducible hsp12 mRNA are shown for the different PQC deletions.
C. Expression of canonical HSR genes (stressed for 6 hours at 37°C) are probed in these PQC deletion strains. The fold change in their expression values in the PQC deletion strains w.r.t By4741 (parental strain) was compared to the global alteration in transcripts using Mann-Whitney test. The p-values obtained were corrected for multiple comparisons using FDR and the cells that did not cross the cut-off significance for p-value 0.05 are without any colour. The colour scheme is graded from red (down-regulated) to blue (upregulated) showing median fold change values. The size of the circle represents the median fold change shift in either direction.

**Figure S3.**
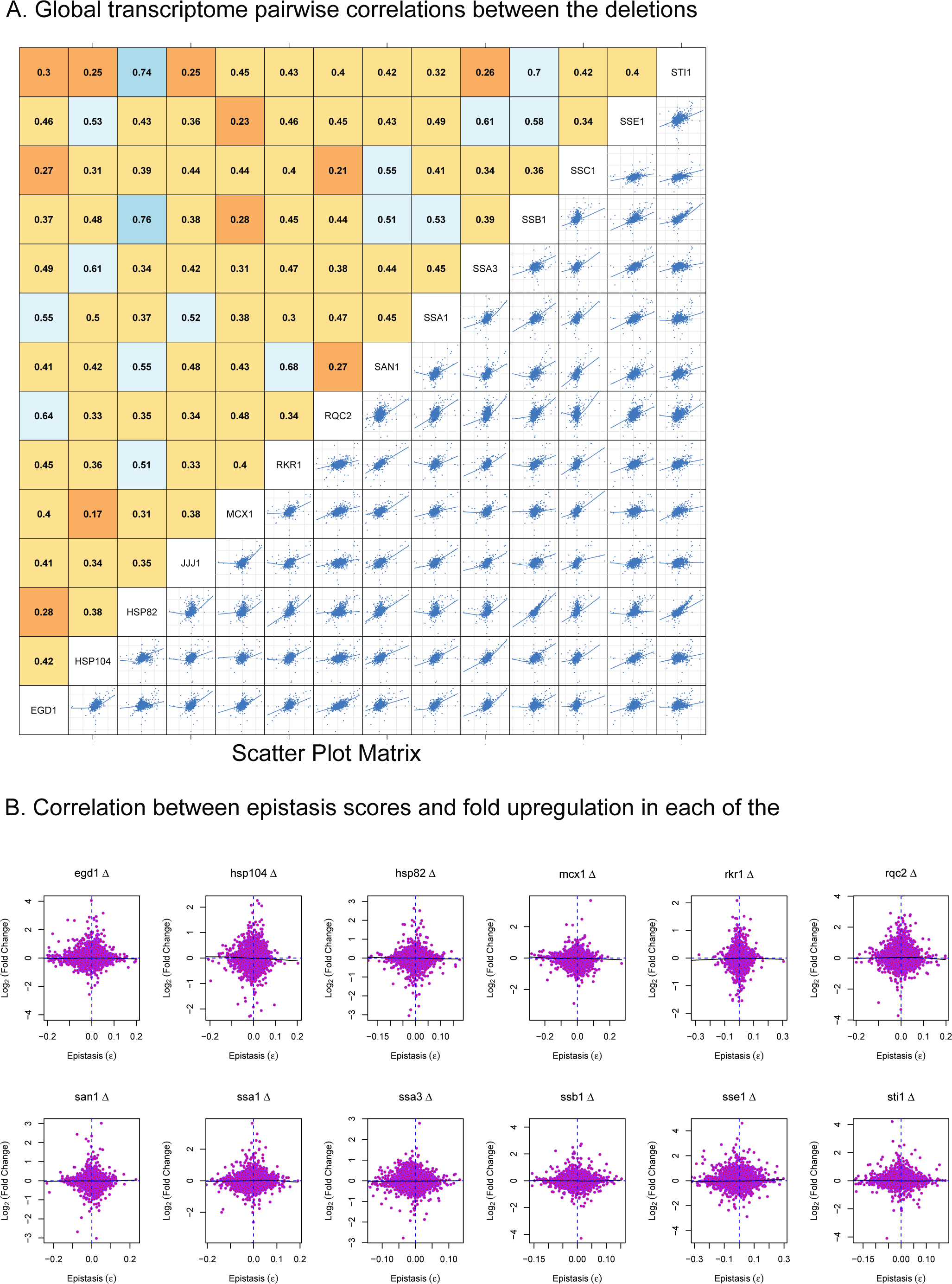
A. For each pair of the PQC deletions transcriptome alteration was quantified w.r.t WT strain. Pairwise comparison was then done between the different PQC deletions using transcriptome alterations. The upper triangular matrix shows the Pearson correlation between the PQC deletions. Lower triangular part of the graph are scatter plots of the transcriptomes of the deletion pairs. The trend lines are LOESS fits and are guides for visualization.
B. Transcriptional alteration of each gene (fold change in a PQC gene deletion with respect to WT strain from this study) is plotted against their Boone epistasis score (ε) as reported (20).

**Figure S4.**
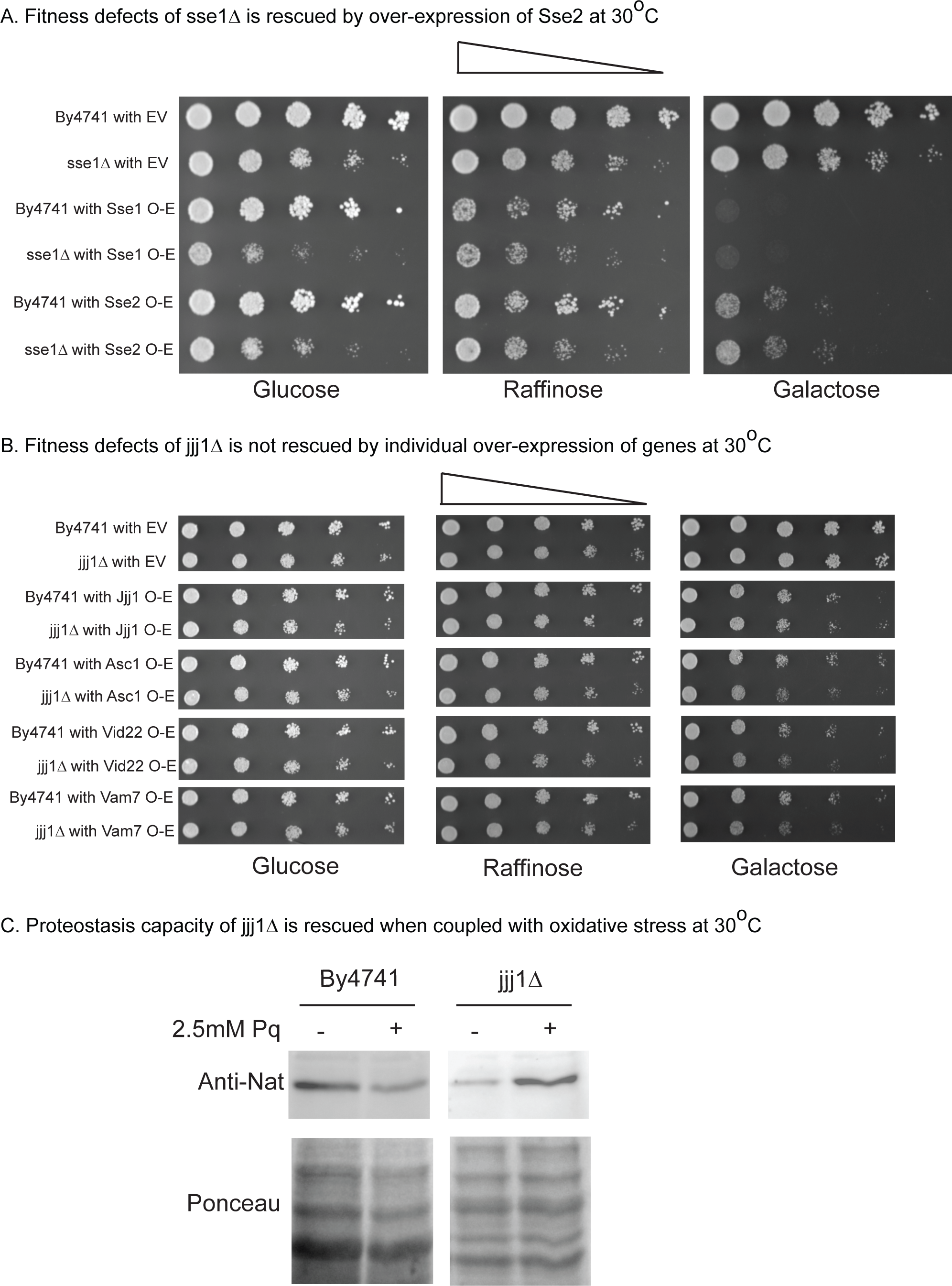
A. By4741 and sse1Δ bearing either empty vector or galactose-inducible over expression of Sse1 or Sse2 were spotted on YPD, YP + Raffinose or YP+ Galactose and grown at 30°C.
B. By4741 and jjj1Δ bearing either empty vector or galactose-inducible over expression of Jjj1 or Asc1 or Vam7 or Vid22 were spotted on YPD, YP + Raffinose or YP+ Galactose and grown at 30°C.
C. Steady state amounts of Wt Nat and TS22 at 30°C in By4741 and jjj1Δ with or without 2.5mM paraquat treatment.

